# Whole genome re-sequencing reveals recent signatures of selection in three strains of farmed Nile tilapia (*Oreochromis niloticus*)

**DOI:** 10.1101/825364

**Authors:** María I. Cádiz, María E. López, Diego Díaz-Domínguez, Giovanna Cáceres, Grazyella M. Yoshida, Daniel Gomez-Uchida, José M. Yáñez

**Affiliations:** Facultad de Ciencias Veterinarias y Pecuarias, Universidad de Chile, Avenida Santa Rosa 11735, 8820808, La Pintana, Santiago, Chile; Programa de Doctorado en Ciencias Silvoagropecuarias y Veterinarias, Campus Sur, Universidad de Chile, Santa Rosa 11315, La Pintana, Santiago, Chile. CP: 8820808; Department of Animal Breeding and Genetics, Swedish University of Agricultural Sciences, Uppsala, Sweden; Departamento de Ciencias de la Computación, Universidad de Chile; Facultad de Ciencias Naturales y Oceanográficas, Universidad de Concepción, Concepción, Chile; Núcleo Milenio INVASAL, Concepción, Chile

**Keywords:** Selection signatures, Nile tilapia, domestication, whole-genome sequencing, *Oreochromis niloticus*, SNPs

## Abstract

Nile tilapia (*Oreochromis niloticus* Linnaeus, 1758) belong to the second most cultivated group of fish in the world, mainly because of its favorable characteristics for production. Genetic improvement programs in this species began in the late 1980s to enhance some traits of commercial interest. The resulting domestication process of Nile tilapia may have modified the genome through selective pressure, leaving signals that can be detected at the molecular level. In this work, signatures of selection were identified using genome-wide SNP data, using two complementary methods based in extended haplotype homozygosity (EHH)._Whole-genome sequencing of 326 individuals from three strains (A, B and C) of farmed tilapia from two countries (Brazil and Costa Rica) was carried out using Illumina HiSeq 2500 technology. After applying conventional SNP-calling and quality-control pipelines, a total of ~1.3M high-quality SNPs were inferred and used as input for the Integrated Haplotype Score (|iHS|) and standardized log-ratio of integrated EHH between pairs of populations (Rsb) methods. We detected 16, 174 and 96 candidate genes subjected to selection in strain A, B, and C, respectively. These candidate genes represent putative genomic landmarks that could contain functions of biological and commercial interest.

## Introduction

Nile tilapia (*Oreochromis niloticus*) is a teleost fish of the Cichlidae family and native to Africa and the Middle East. The geographic range of the species extends from 8ºN to 32ºN ^1,2^. The first record of domestication is dated around 3500 years ago as evidenced in paintings at the Theban tombs in Egypt ^3^. Nowadays, this species is the second most cultivated group of aquaculture species in the world ^4^. Favorable characteristics for production include, but are not limited to, rapid growth, adaptability to different culture conditions, tolerance to high densities, disease resistant, easy reproduction, and tolerance to low concentrations of oxygen ^5^.

Genetic improvement programs (GIPs) for Nile tilapia began in 1988 as a method to handle the production decrease generated by introgressions with Mozambique tilapia (*Oreochromis mossambicus*) ^6,7^. Since then, nearly twenty GIPs have been established for Nile tilapia around the world ^8,9^. GIPs aim to improve traits of commercial interest, such as growth rate, disease resistance and cold and salinity tolerances ^8^. The first Nile tilapia strain created was GIFT (Genetic Improvement of Farmed Tilapia). GIFT was developed by the ICLARM (International Centre for Living Aquatic Resources Management) now the WorldFish Center, in collaboration with the Norwegian Institute of Aquaculture Research (AKVAFORSK, now NOFIMA Marin) and other institutions ^2^. GIFT is a success case for the implementation of GIPs because growth rate in Nile tilapia has doubled in five generations, showing that this species had a positive response to selection ^2^.

Domestication is the process of constant evolutionary and genetic changes in response to captivity ^10^. Nile tilapia can be considered to have reached the level of true domestication (level 5), according to the five categories of the domestication process, because selective breeding programmes have focused on specific goals ^11,12^. This process may have shaped the genetic diversity of Nile tilapia, leaving signatures in their genomes that can be traced. These signatures can: (i) exhibit increased allele frequencies in favorable adaptive substitutions ^13,14^, (ii) show strong linkage disequilibrium (LD) in areas surrounding the signature, which decays downstream and upstream of this region ^15^, and (iii) undergo loss of genetic diversity (selective sweep) and the rate of “effective recombination” in the genome of domestic species compared to the genomes of wild relatives ^16^.

Selection signatures can be detected by scanning the genome of sampled individuals of a given population to search for deviations in allele frequency spectrum (Tajima’s D and Fay and Wu’s H scores), higher or lower population differentiation than under neutral expectations (Fst value) or based on measures of LD (EHH, iHS, Rsb methods) (Vitti, Grossman, and Sabeti 2013). The most suitable method to detect selection signatures depends on the number of populations under study, temporal context scale, and type of selection signatures ^17,18^. Thus, more than one approach is often required to capture any signal in the genome ^19^. For example, methods derived from EHH are used to detect recent positive selection within-population (iHS) and between-populations (Rsb) ^20^, whereas methods based on Fst are expected to identify older selection events ^21^ between-populations^22^.

Several studies of selection signatures have been carried out in aquaculture species ^23–27^. Among tilapia and related species, there are only two studies on selection signatures; one in cichlid fish ^28^ and another describing whole-genome selection signatures in tilapia ^29^. The purpose of the present study was to identify recent signatures of selection in three domestic populations of Nile tilapia from Brazil (A strain) and Costa Rica (B and C strains). We used whole-genome sequencing data and applied two statistical approaches to identify genomic regions putatively under selection: (i) Integrated Haplotype Score, iHS and (ii) standardized log-ratio of integrated EHH (iES) between pairs of populations (Rsb). Finally, the genes under selection were associated to biological functions by performing an enrichment analysis. For this purpose, we used Gene Ontology (GO) and Kyoto Encyclopedia of Genes and Genomes (KEGG) pathways terms.

## Results

### Quality control

Approximately 76.6M raw reads (SD=65.0 million reads) per fish were generated for the 326 individuals through whole genome re-sequencing. From these, 99.6% were successfully mapped to the reference genome of Nile tilapia. The mean read coverage per individual was 8.7X (SD=9.9X). Subsequent variant calling yielded a total of 38.45M variants discovered. From this set, only 1.3M variants were shared among all three populations and 280 individuals were kept after quality control, which were used for the following analysis (23 individuals with call rate below 80% and 23 with IBD>0.5 were removed; Table 2). Of this set of SNPs, based in the results from SnpEff: 30.3% of the SNPs were located within intron regions, 34.56% transcript, 12.6% downstream, 12.4% upstream, 5.2% in intergenic regions and 3.1% in exon regions. The exon region contained 56.7% silent mutations and 42.9% missense mutations. The missense to silent ratio was 0.76%.

### Basic statistics and population structure analysis

Observed and expected heterozygosity (H_o_/H_e_) obtained were 0.25/0.31, 0.25/0.30 and 0.23/0.30 for A, B, and C strains, respectively (Table 1). All these genetic diversity measures were statistically significant (p < 0.05, Kruskal–Wallis test). While the Weir and Cockerham mean (Fst) values among the three strains were low and very similar: A vs B = 0.045 (CI = 0.0445-0.0446), A vs C = 0.045 (CI = 0.0446-0.0449), and B vs C = 0.042 (CI = 0.0413-0.0416).

**Table 1.** Description of strains of Nile tilapia analyzed in this study.

| *Strain code* | *GeO*<sup>1</sup> | *n*<sup>2</sup> | *Ho*<sup>3</sup> | *Sd*<sup>4</sup> | *IC*<sup>5</sup> | *He*<sup>6</sup> | *Sd*<sup>7</sup> | *IC*<sup>7</sup> |
| --- | --- | --- | --- | --- | --- | --- | --- | --- |
| A | Brazil | 56 | 0.236 | 0.119 | 0.236-0.237 | 0.306 | 0.126 | 0.306-0.306 |
| B | Costa Rica | 100 | 0.253 | 0.121 | 0.253-0.254 | 0.298 | 0.124 | 0.298-0.298 |
| C | Costa Rica | 124 | 0.233 | 0.112 | 0.232-0.233 | 0.299 | 0.124 | 0.299-0.299 |

<sup>1</sup> *GeO*, Geographical origin; <sup>2</sup> *n*, number of samples; <sup>3</sup> *H<sub>o</sub>*, observed heterozygosity; <sup>4</sup> *Sd*, Standard deviation of *H<sub>o</sub>*; <sup>5</sup> *IC*, confidence interval of *H<sub>o</sub>*; <sup>6</sup> *H<sub>e</sub>*, expected heterozygosity; <sup>7</sup> *Sd*, Standard deviation of *H<sub>e</sub>*; <sup>8</sup> *IC*, confidence interval of *H<sub>e</sub>*.

**Table 2.** Quality control (QC) of SNPs and individuals.

| *QC*<sup>1</sup> | *A*<sup>2</sup> | *B*<sup>3</sup> | *C*<sup>4</sup> | *Match/total*<sup>5</sup> |
| --- | --- | --- | --- | --- |
| ***SNPs*** |  |  |  |  |
| Initial number of SNPs | 36.399.577 |  |  |  |
| Pred score<30 | 22.039.340 |  |  |  |
| Call rate/SNPs 90% | 9.533.967 |  |  |  |
| HWE(1x10-9) | 9.532.139 | 9.520.991 | 9.498.112 |  |
| MAF (0.05) | 2.721.234 | 3.170.541 | 2.888.496 | 1.333.203<sup>5</sup> |
| ***Individuals*** |  |  |  |  |
| Initial N° of individuals | 61 | 126 | 139 | 326 |
| Call rate 80% | 59 | 113 | 131 | 303 |
| IBD | 56 | 100 | 124 | 280<sup>8</sup> |

<sup>1</sup> *QC*, quality control; strains <sup>2</sup>*A*, <sup>3</sup>*B* and <sup>4</sup>*C*; <sup>5</sup> *Match/total*, match SNPs among strains and total of individuals; <sup>6</sup> 1.333.203, match of SNPs among strains using analysis of signatures of selection (iHS and Rsb), <sup>5</sup> 1.333.203, match of SNPs among strains using population analysis; <sup>6</sup> 280, number of individuals in all analysis.

Regarding the genetic structure, the principal component analysis (PCA) (Figure 1) shows three distinct clusters corresponding to strain A, B and C of Nile tilapia. The first two eigenvectors together explain of 22.45% of the genetic variability. Using the first principal component (PC1), the first two clusters correspond to strains A and B, and the third one corresponds to strain C. In addition, admixture analysis revealed that the best K=7 (Figure 2), is in agreement with the expected level of admixture for the strains studied here.

**Figure 1.**
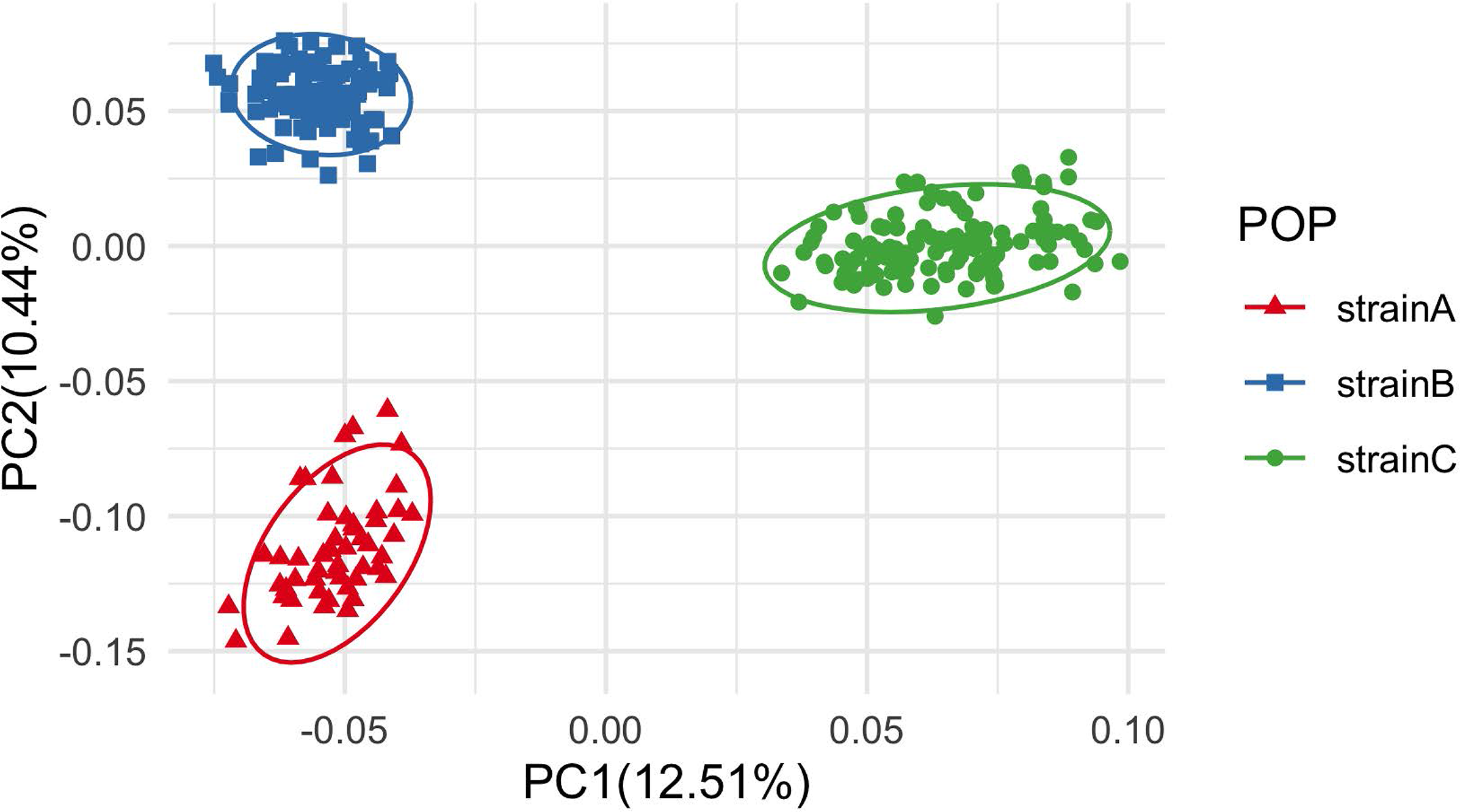
Principal component analysis (PCA) of genetic differentiation among 280 individuals based on ~1.3M SNPs. Each dot represents one individual. Strain A (red triangle); Strain B (blue rectangle) and Strain C (green circle).

**Figure 2.**
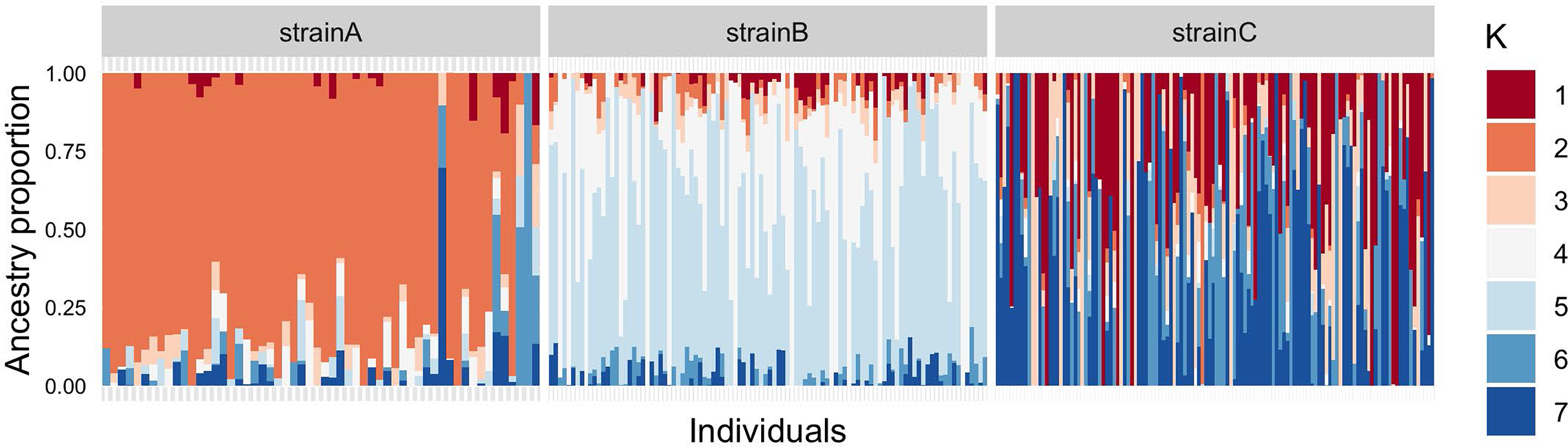
Admixture analysis of K=7 for Nile tilapia populations: strains A; strain B and strain C. Each color represents a different theoretical ancestral population and each individual is represented by a vertical bar.

### Signatures of selection

The *iHS* analysis revealed signatures of selection in the three strains studied (Figure 3; Supplementary Table S2). Strain A showed 17 SNPs surpassing the significance threshold, with 310 genes localized within the 1Mb windows of each marker in linkage group (LG) 3 and 9 (Table S2-A). Strain B showed 81 SNPs surpassing the significance threshold, with 944 genes localized within the 1Mb windows of each marker in seven different LG (2, 3, 10, 13, 14, 15 and 18) (Supplementary Table S2-B). Finally, strain C showed 34 SNPs surpassing the significance threshold, with 734 genes localized within the 1Mb window of each marker in six different LG (2, 3, 9, 10, 13 and 15) (Supplementary Table S2-C).

**Figure 3.**
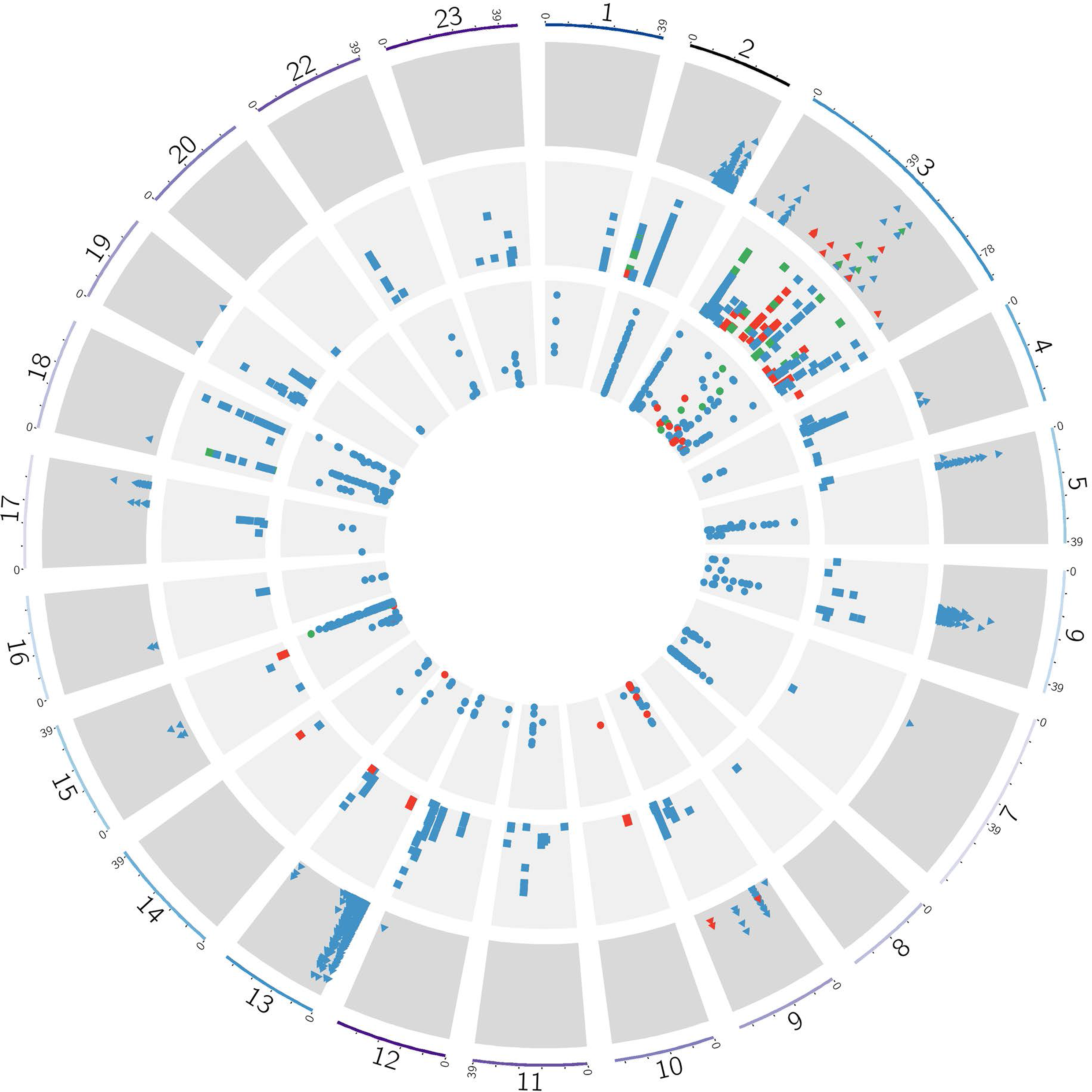
Nile tilapia genome showing signatures of selection in strains A, B, and C: strain A, triangle; strain B, rectangle and strain C, circle. The radial plot illustrates the distribution of iHS and Rbs scores across the genome of Nile tilapia: iHS scores (red); Rsb (blue) and matched SNPs between methods (green). The final radial plot corresponds to the chromosomes.

The *Rsb* method detected several signatures of selection between strains (Figure 3, Supplementary Table S3). The strain A-B comparison identified 1394 SNPs surpassing the significance threshold, with 980 SNPs showing evidence of selection in strain A and 414 SNPs showing evidence of selection in strain B. The strain B-C comparison identified 839 SNPs surpassing the significance threshold, with 323 SNPs showing evidence of selection in strain B and 516 SNPs showing evidence of selection in strain C. Finally, the strain C-A comparison identified 1167 SNPs surpassing the significance the threshold, with 295 SNPs showing evidence of selection in strain C and 872 SNPs showing evidence of selection in strain A.

For strain A 1288 SNPs with unique positions showed evidence of selection, which were associated with 1403 genes localized in fourteen LG (excluding 1, 8, 10, 11, 14, 20, 22 and 23) (Supplementary Table S3-A). In strain B strain 622 SNPs showed evidence of selection, which were associated with 2613 genes distributed in all LG (Supplementary Table S3-B). And for strain C 649 SNPs showed evidence of selection, which were associated with 2023 genes localized in nineteen LG (all chromosomes, excluding chromosomes 8 and 19) (Supplementary Table S3-C).

*iHS and Rsb methods*. As mentioned above strong candidate regions for selection were defined as those genomic regions containing SNPs with values above the significance threshold detected by both methods (Figure 3). For each strain we detected several candidate genes subjected to selection within 1Mb windows containing SNPs surpassing the significance threshold by both methods. For strain A we detected 16 genes localized in LG 3 and 9. For strain B we detected 174 genes, localized in seven different LG (2, 3, 10, 13, 14,15 and 18). For strain C we detected 96 genes, localized in five different LG (2, 3, 9, 13 and 15). Four genes were detected by both methods across all strains (A, B and C): ribonuclease inhibitor-like, BTN1A1, MOG and Ladderlectin localized in chromosome 3. In addition, 57 genes were detected by both methods and shared across two strains from the same geographical origin (B and C). In contrast, a lower number of genes were common between strain A from Brazil and both strains B and C from Costa Rica (AB = eight genes, AC = five genes).

### Functional enrichment analysis

The results of enrichment analysis of the total signals of selection detected by both *iHS* and *Rsb* methods are shown in Supplementary Table S4.

For strain A, we detected 1483 proteins contained within the 1Mb windows of all SNPs identified under selection. From these sequences, 1185 were functionally annotated via BLAST (zebrafish) and 1147 genes were detected by DAVID software (Table 3; Figure 4; Supplementary Table S4-A). By using gene ontology (GO): we determined that 58.8% of the genes (n = 675) were classified within the category Biological Process (BP), with 33 enriched terms, of which 16 were significant; 60.7% of the genes (n = 696) were classified within category Cellular Component (CC), with 10 enriched terms, of which 8 were significant; and 58.8% of the genes (n = 674) were classified within the category Molecular Function (MF), with five enriched terms, of which three were significant. For category KEGG pathway we detected 25.8% of genes (n = 296), with six enriched terms, of which four were significant.

**Table 3.** Enriched GO and KEGG pathways term for genes in regions under selection of strain A of Nile tilapia related to domestication.

| *Code<sup>1</sup>* | *Term<sup>2</sup>* | *Genes<sup>3</sup>* | *P-value<sup>4</sup>* |
| --- | --- | --- | --- |
| ***Biological Process 58.8% (675)*** |  |  |  |
| GO:0050890 | cognition <sup>(B)</sup> | 5 | 0.004* |
| GO:0002040 | sprouting angiogenesis <sup>(R)</sup> | 10 | 0.018* |
| GO:0007612 | learning <sup>(B)</sup> | 4 | 0.023* |
| GO:0060537 | muscle tissue development <sup>(G)</sup> | 15 | 0.032* |
| GO:0048513 | animal organ development <sup>(G)</sup> | 116 | 0.042* |
| GO:0090594 | inflammatory response to wounding <sup>(I)</sup> | 3 | 0.042* |
| GO:0001501 | skeletal system development <sup>(G)</sup> | 21 | 0.044* |
| GO:0003171 | atrioventricular valve development <sup>(G)</sup> | 4 | 0.047* |
| GO:0035265 | organ growth <sup>(G)</sup> | 5 | 0.050* |
| GO:0006936 | muscle contraction <sup>(G)</sup> | 9 | 0.072 |
| GO:0003170 | heart valve development <sup>(G)</sup> | 4 | 0.080 |
| GO:0048514 | blood vessel morphogenesis <sup>(G)</sup> | 23 | 0.086 |
| GO:0061448 | connective tissue development <sup>(G)</sup> | 11 | 0.088 |
| GO:0090257 | regulation of muscle system process <sup>(G)</sup> | 5 | 0.089 |
| GO:0060536 | cartilage morphogenesis <sup>(G)</sup> | 4 | 0.089 |
| GO:1902850 | microtubule cytoskeleton organization involved in mitosis <sup>(G)</sup> | 4 | 0.089 |
| GO:0007018 | microtubule-based movement <sup>(G)</sup> | 13 | 0.091 |
| GO:0008038 | neuron recognition <sup>(B)</sup> | 4 | 0.099 |
| ***Cellular Component 60.7% (696)*** |  |  |  |
| GO:0005884 | actin filament <sup>(G)</sup> | 6 | 0.008* |
| GO:0031941 | filamentous actin <sup>(G)</sup> | 4 | 0.017* |
| GO:0031674 | I band <sup>(G)</sup> | 5 | 0.096 |
| ***Molecular Function 58.8% (674)*** |  |  |  |
| GO:0015631 | Tubulin binding <sup>(G)</sup> | 15 | 0.036* |

| *KEGG 25.8% (296)* |  |  |  |
| --- | --- | --- | --- |
| dre04114 | Oocyte meiosis <sup>(R)</sup> | 15 | 0.017* |
| dre04261 | Adrenergic signaling in cardiomyocytes <sup>(G)</sup> | 18 | 0.026* |
| dre04310 | Wnt signaling pathway <sup>(G)</sup> | 15 | 0.074 |

<sup>1</sup> Code associated to GO and KEGG pathways terms; <sup>2</sup> Term enriched; <sup>3</sup> Number of genes enriched associated with term; <sup>4</sup> P-value, significance. Traits associated with domestication: <sup>(G)</sup>Growth; <sup>(B)</sup>Behavior; <sup>(I)</sup>Immune system; <sup>(R)</sup>Reproduction; <sup>(A)</sup>Adaptation.

**Figure 4.**
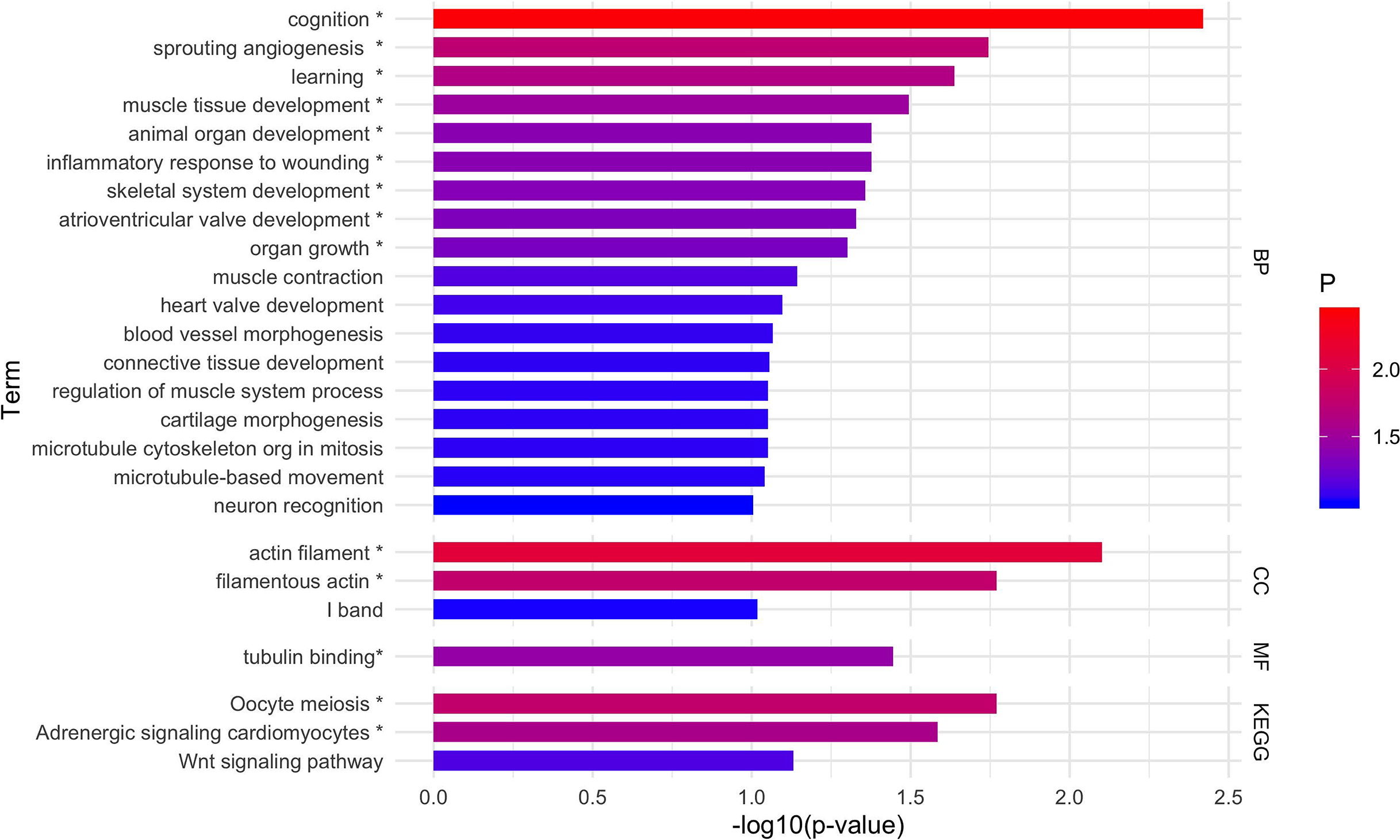
Enrichment analysis for GO and KEGG pathways term for strain A by DAVID. Each bar represents the -log10(p-value) for term. The colors represent the significance of each term; red color presents significant values and as the significance decreases the gradient turns blue. * significant terms.

For strain B, we detected 3074 proteins contained within the 1Mb windows of all SNPs identified under selection. From these sequences, 1836 were functionally annotated via BLAST (zebrafish) and 1800 genes were detected for DAVID software (Table 4; Figure 5; Supplementary Table S4-B). By using GO: we determined that 58.6% of the genes (n = 1055) were classified within the category BP, with 43 enriched terms, of which 29 were significant; 55.4% of the genes (n = 977) were classified within the category CC, with 19 enriched terms, of which 14 were significant; and 57.6% of the genes (n = 1036) were classified within the category MF, with 13 enriched terms, of which 10 were significant. For category KEGG pathway we detected 24.3% of the genes (n = 434), with 15 enriched terms, of which 10 were significant.

**Table 4.**
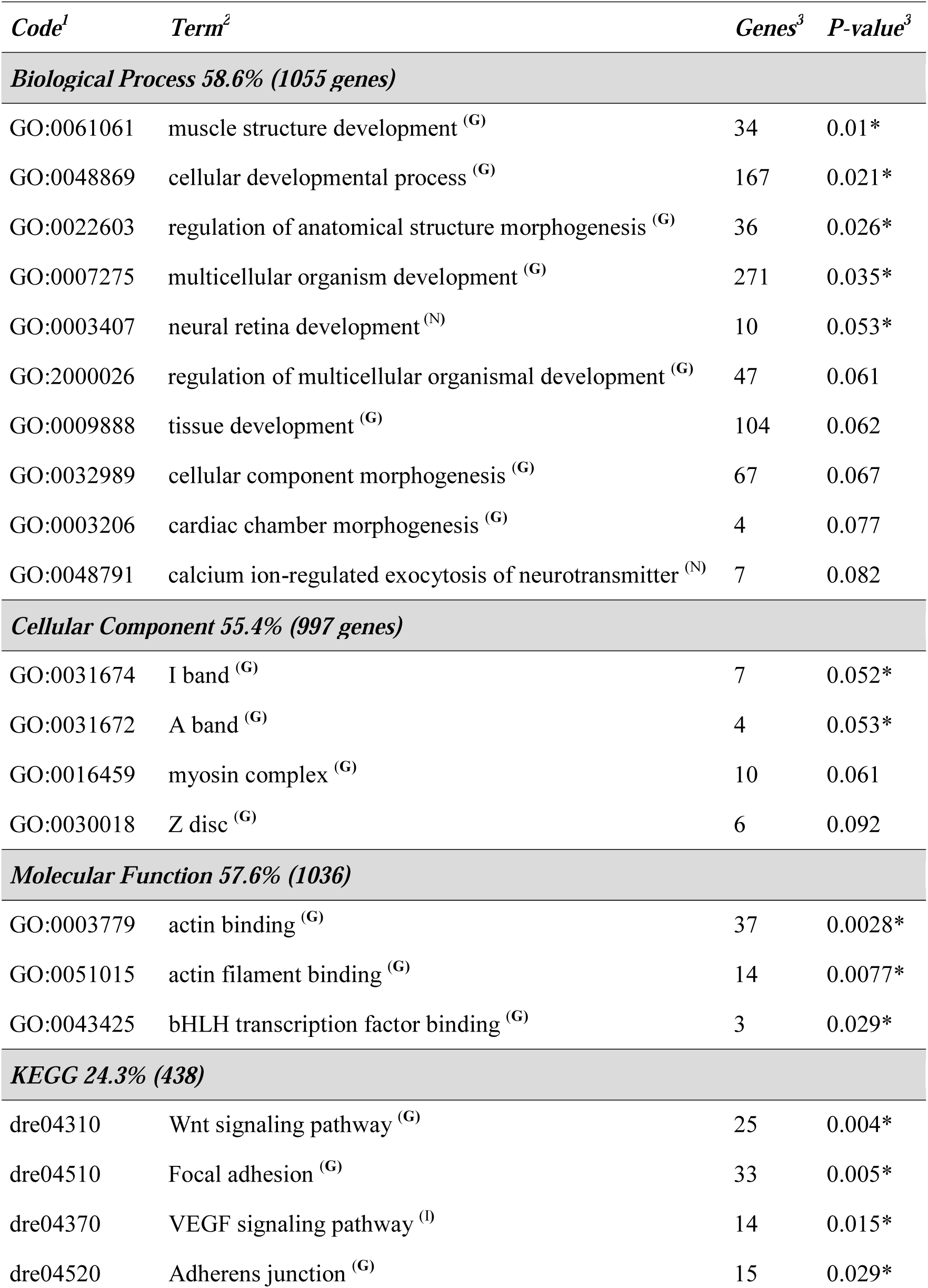
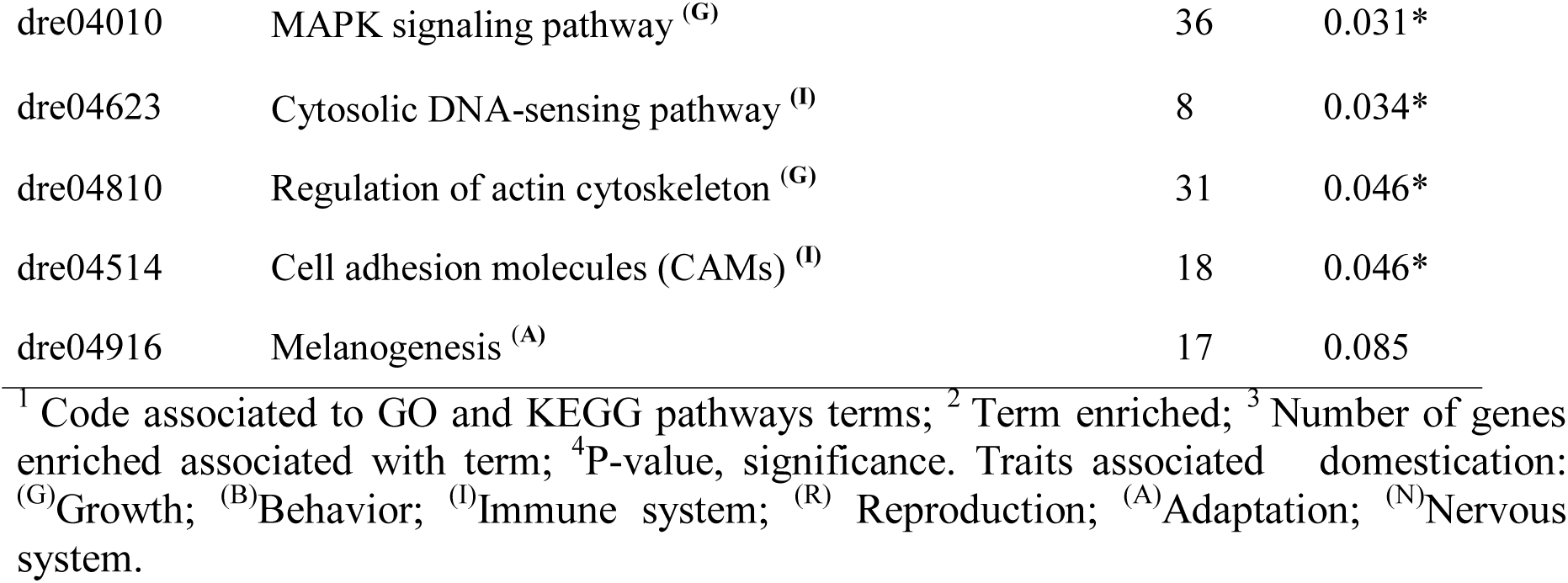
Enriched GO and KEGG pathways term for genes in regions under selection of strain B of Nile tilapia related to domestication.

| *Code<sup>1</sup>* | *Term<sup>2</sup>* | *Genes<sup>3</sup>* | *P-value<sup>3</sup>* |
| --- | --- | --- | --- |
| **Biological Process 58.6% (1055 genes)** |  |  |  |
| GO:0061061 | muscle structure development <sup>(G)</sup> | 34 | 0.01* |
| GO:0048869 | cellular developmental process <sup>(G)</sup> | 167 | 0.021* |
| GO:0022603 | regulation of anatomical structure morphogenesis <sup>(G)</sup> | 36 | 0.026* |
| GO:0007275 | multicellular organism development <sup>(G)</sup> | 271 | 0.035* |
| GO:0003407 | neural retina development <sup>(N)</sup> | 10 | 0.053* |
| GO:2000026 | regulation of multicellular organismal development <sup>(G)</sup> | 47 | 0.061 |
| GO:0009888 | tissue development <sup>(G)</sup> | 104 | 0.062 |
| GO:0032989 | cellular component morphogenesis <sup>(G)</sup> | 67 | 0.067 |
| GO:0003206 | cardiac chamber morphogenesis <sup>(G)</sup> | 4 | 0.077 |
| GO:0048791 | calcium ion-regulated exocytosis of neurotransmitter <sup>(N)</sup> | 7 | 0.082 |
| **Cellular Component 55.4% (997 genes)** |  |  |  |
| GO:0031674 | I band <sup>(G)</sup> | 7 | 0.052* |
| GO:0031672 | A band <sup>(G)</sup> | 4 | 0.053* |
| GO:0016459 | myosin complex <sup>(G)</sup> | 10 | 0.061 |
| GO:0030018 | Z disc <sup>(G)</sup> | 6 | 0.092 |
| **Molecular Function 57.6% (1036)** |  |  |  |
| GO:0003779 | actin binding <sup>(G)</sup> | 37 | 0.0028* |
| GO:0051015 | actin filament binding <sup>(G)</sup> | 14 | 0.0077* |
| GO:0043425 | bHLH transcription factor binding <sup>(G)</sup> | 3 | 0.029* |
| **KEGG 24.3% (438)** |  |  |  |
| dre04310 | Wnt signaling pathway <sup>(G)</sup> | 25 | 0.004* |
| dre04510 | Focal adhesion <sup>(G)</sup> | 33 | 0.005* |
| dre04370 | VEGF signaling pathway <sup>(I)</sup> | 14 | 0.015* |
| dre04520 | Adherens junction <sup>(G)</sup> | 15 | 0.029* |
| dre04010 | MAPK signaling pathway <sup>(G)</sup> | 36 | 0.031* |
| dre04623 | Cytosolic DNA-sensing pathway <sup>(I)</sup> | 8 | 0.034* |
| dre04810 | Regulation of actin cytoskeleton <sup>(G)</sup> | 31 | 0.046* |
| dre04514 | Cell adhesion molecules (CAMs) <sup>(I)</sup> | 18 | 0.046* |
| dre04916 | Melanogenesis <sup>(A)</sup> | 17 | 0.085 |

<sup>1</sup> Code associated to GO and KEGG pathways terms; <sup>2</sup> Term enriched; <sup>3</sup> Number of genes enriched associated with term; <sup>4</sup> P-value, significance. Traits associated domestication: <sup>(G)</sup>Growth; <sup>(B)</sup>Behavior; <sup>(I)</sup>Immune system; <sup>(R)</sup> Reproduction; <sup>(A)</sup>Adaptation; <sup>(N)</sup>Nervous system.

**Figure 5.**
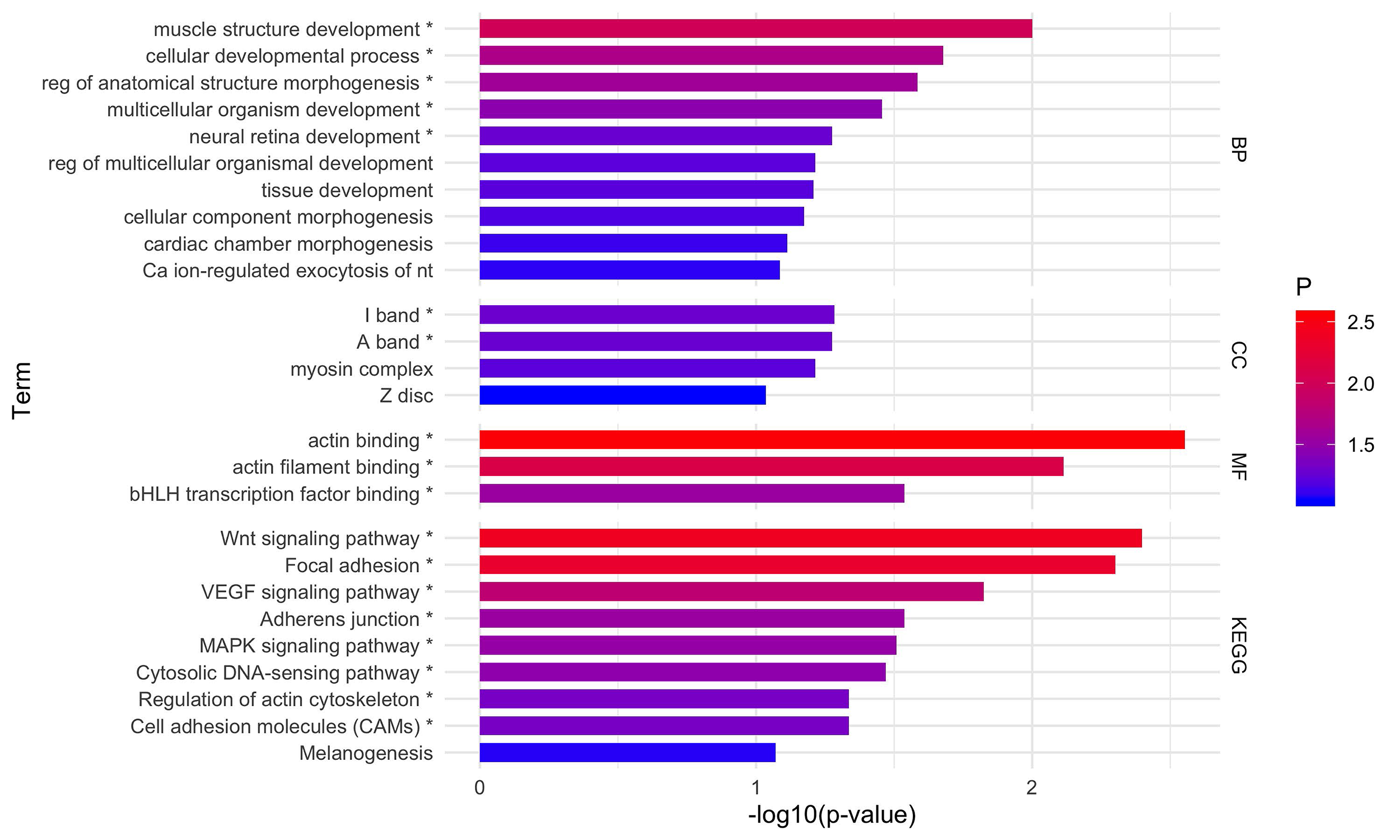
Enrichment analysis for GO and KEGG pathways term for strain B by DAVID. Each bar represents the -log10(p-value) for term. The colors represent the significance of each term; red color presents significant values and as the significance decreases the gradient turns blue. * significative terms.

Finally, for strain C we detected 2139 proteins contained within the 1Mb windows of all SNPs identified under selection. From these sequences, 1456 were functionally annotated via BLAST (zebrafish) and 1400 genes were detected for DAVID software (Table 5; Figure 6; Supplementary Table S4-C). By using GO: we determined that 59% of the genes (n = 826) were classified within the category BP, with 34 enriched terms, of which 21 were significant; 59.9% of the genes (n = 839) were classified within the category CC, with nine enriched terms, of which four were significant; and 44.9% of the genes (n = 628) were classified within the category MF, with 17 enriched terms, of which six were significant. For category KEGG pathway we detected 22.9% of the genes (n = 320), with seven enriched terms, of which four were significant.

**Table 5.**
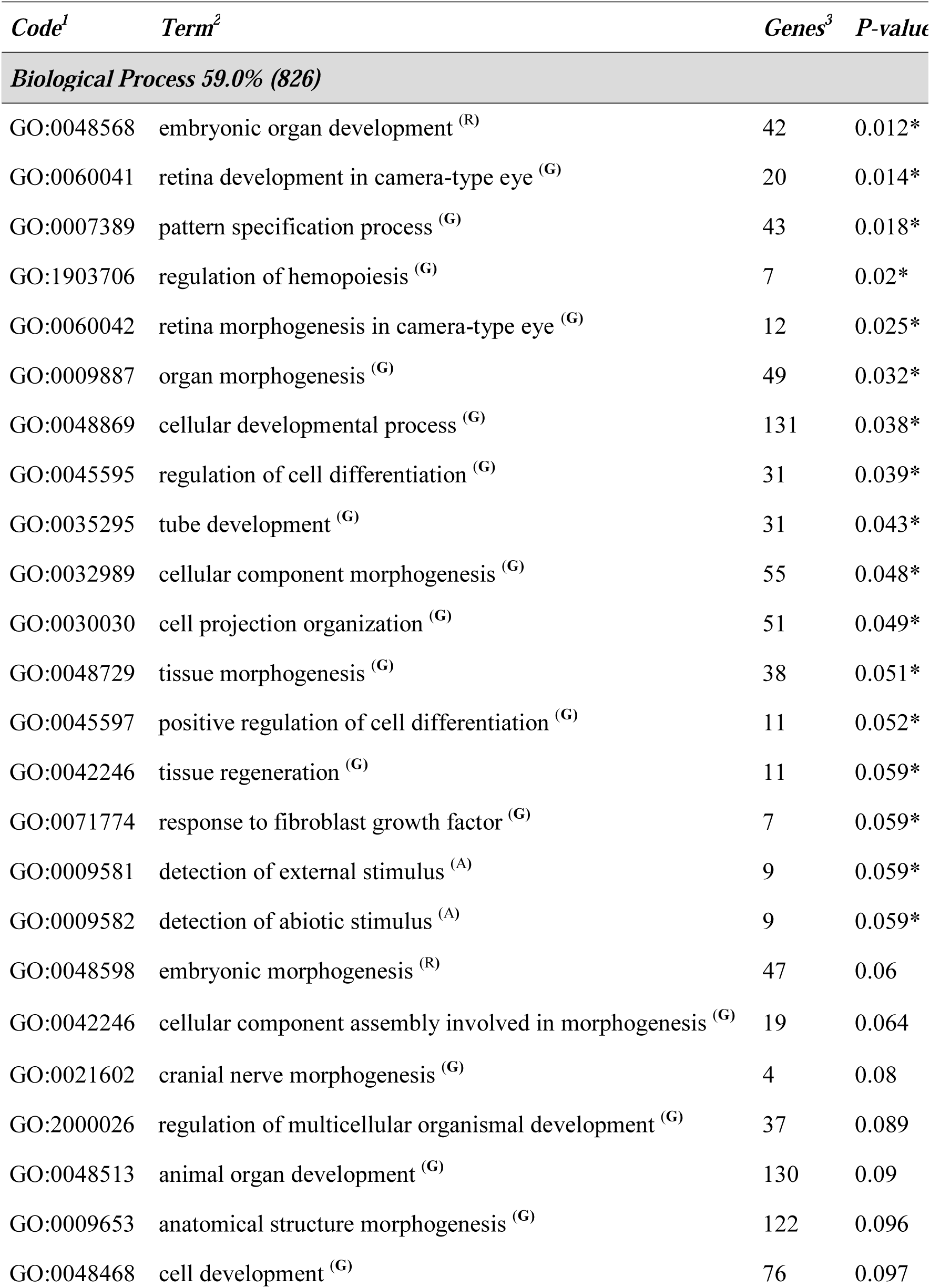
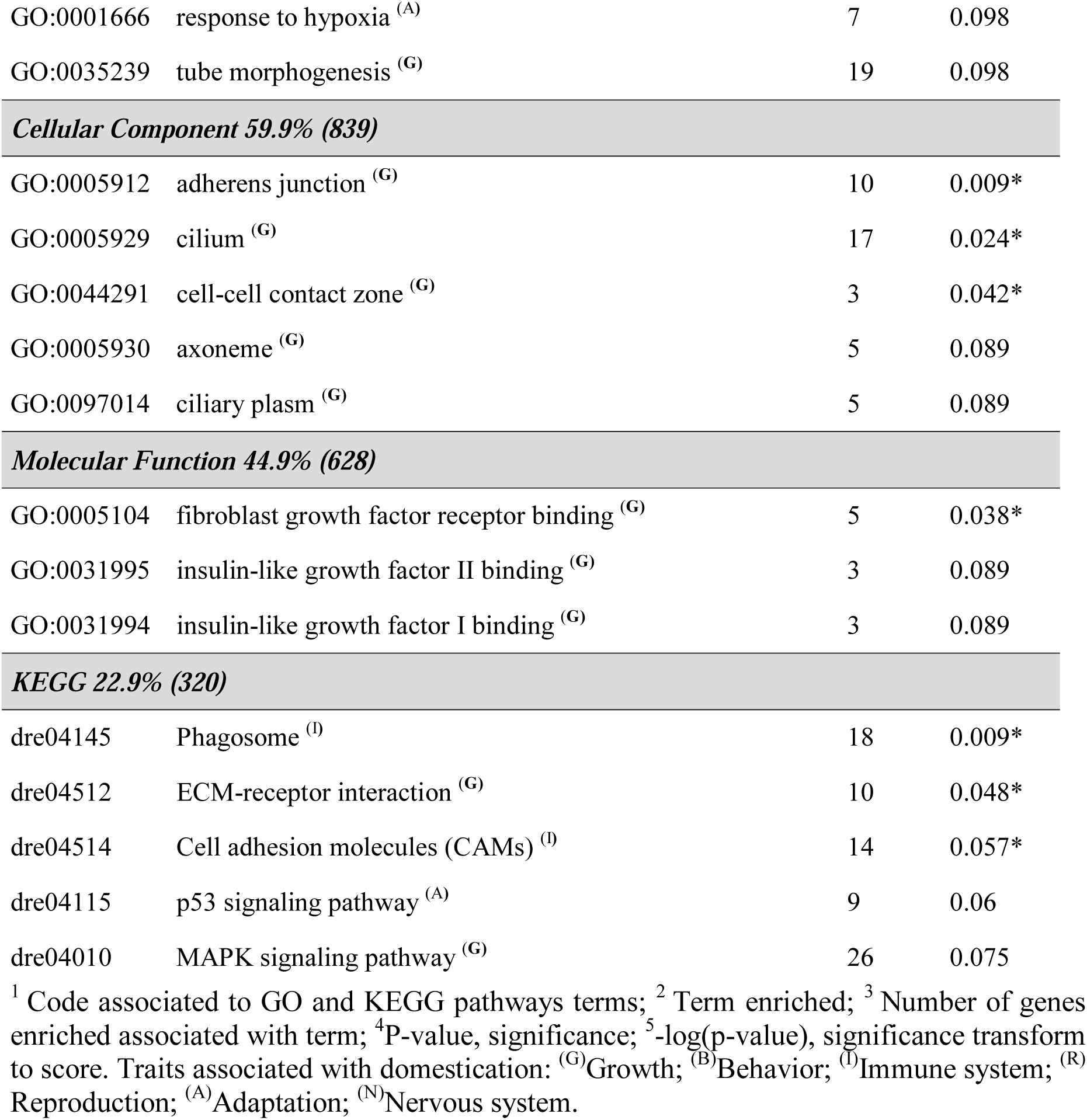
Enriched GO and KEGG pathways term for genes in regions under selection of strain C of Nile tilapia related to domestication.

| *Code*<sup>1</sup> | *Term*<sup>2</sup> | *Genes*<sup>3</sup> | *P-value*<sup>4</sup> |
| --- | --- | --- | --- |
| **Biological Process 59.0% (826)** |  |  |  |
| GO:0048568 | embryonic organ development <sup>(R)</sup> | 42 | 0.012* |
| GO:0060041 | retina development in camera-type eye <sup>(G)</sup> | 20 | 0.014* |
| GO:0007389 | pattern specification process <sup>(G)</sup> | 43 | 0.018* |
| GO:1903706 | regulation of hemopoiesis <sup>(G)</sup> | 7 | 0.02* |
| GO:0060042 | retina morphogenesis in camera-type eye <sup>(G)</sup> | 12 | 0.025* |
| GO:0009887 | organ morphogenesis <sup>(G)</sup> | 49 | 0.032* |
| GO:0048869 | cellular developmental process <sup>(G)</sup> | 131 | 0.038* |
| GO:0045595 | regulation of cell differentiation <sup>(G)</sup> | 31 | 0.039* |
| GO:0035295 | tube development <sup>(G)</sup> | 31 | 0.043* |
| GO:0032989 | cellular component morphogenesis <sup>(G)</sup> | 55 | 0.048* |
| GO:0030030 | cell projection organization <sup>(G)</sup> | 51 | 0.049* |
| GO:0048729 | tissue morphogenesis <sup>(G)</sup> | 38 | 0.051* |
| GO:0045597 | positive regulation of cell differentiation <sup>(G)</sup> | 11 | 0.052* |
| GO:0042246 | tissue regeneration <sup>(G)</sup> | 11 | 0.059* |
| GO:0071774 | response to fibroblast growth factor <sup>(G)</sup> | 7 | 0.059* |
| GO:0009581 | detection of external stimulus <sup>(A)</sup> | 9 | 0.059* |
| GO:0009582 | detection of abiotic stimulus <sup>(A)</sup> | 9 | 0.059* |
| GO:0048598 | embryonic morphogenesis <sup>(R)</sup> | 47 | 0.06 |
| GO:0042246 | cellular component assembly involved in morphogenesis <sup>(G)</sup> | 19 | 0.064 |
| GO:0021602 | cranial nerve morphogenesis <sup>(G)</sup> | 4 | 0.08 |
| GO:2000026 | regulation of multicellular organismal development <sup>(G)</sup> | 37 | 0.089 |
| GO:0048513 | animal organ development <sup>(G)</sup> | 130 | 0.09 |
| GO:0009653 | anatomical structure morphogenesis <sup>(G)</sup> | 122 | 0.096 |
| GO:0048468 | cell development <sup>(G)</sup> | 76 | 0.097 |
| GO:0001666 | response to hypoxia <sup>(A)</sup> | 7 | 0.098 |
| GO:0035239 | tube morphogenesis <sup>(G)</sup> | 19 | 0.098 |
| **Cellular Component 59.9% (839)** |  |  |  |
| GO:0005912 | adherens junction <sup>(G)</sup> | 10 | 0.009* |
| GO:0005929 | cilium <sup>(G)</sup> | 17 | 0.024* |
| GO:0044291 | cell-cell contact zone <sup>(G)</sup> | 3 | 0.042* |
| GO:0005930 | axoneme <sup>(G)</sup> | 5 | 0.089 |
| GO:0097014 | ciliary plasm <sup>(G)</sup> | 5 | 0.089 |
| **Molecular Function 44.9% (628)** |  |  |  |
| GO:0005104 | fibroblast growth factor receptor binding <sup>(G)</sup> | 5 | 0.038* |
| GO:0031995 | insulin-like growth factor II binding <sup>(G)</sup> | 3 | 0.089 |
| GO:0031994 | insulin-like growth factor I binding <sup>(G)</sup> | 3 | 0.089 |
| **KEGG 22.9% (320)** |  |  |  |
| dre04145 | Phagosome <sup>(I)</sup> | 18 | 0.009* |
| dre04512 | ECM-receptor interaction <sup>(G)</sup> | 10 | 0.048* |
| dre04514 | Cell adhesion molecules (CAMs) <sup>(I)</sup> | 14 | 0.057* |
| dre04115 | p53 signaling pathway <sup>(A)</sup> | 9 | 0.06 |
| dre04010 | MAPK signaling pathway <sup>(G)</sup> | 26 | 0.075 |

<sup>1</sup> Code associated to GO and KEGG pathways terms; <sup>2</sup> Term enriched; <sup>3</sup> Number of genes enriched associated with term; <sup>4</sup> P-value, significance; <sup>5</sup> -log(p-value), significance transform to score. Traits associated with domestication: <sup>(G)</sup>Growth; <sup>(B)</sup>Behavior; <sup>(I)</sup>Immune system; <sup>(R)</sup>Reproduction; <sup>(A)</sup>Adaptation; <sup>(N)</sup>Nervous system.

**Figure 6.**
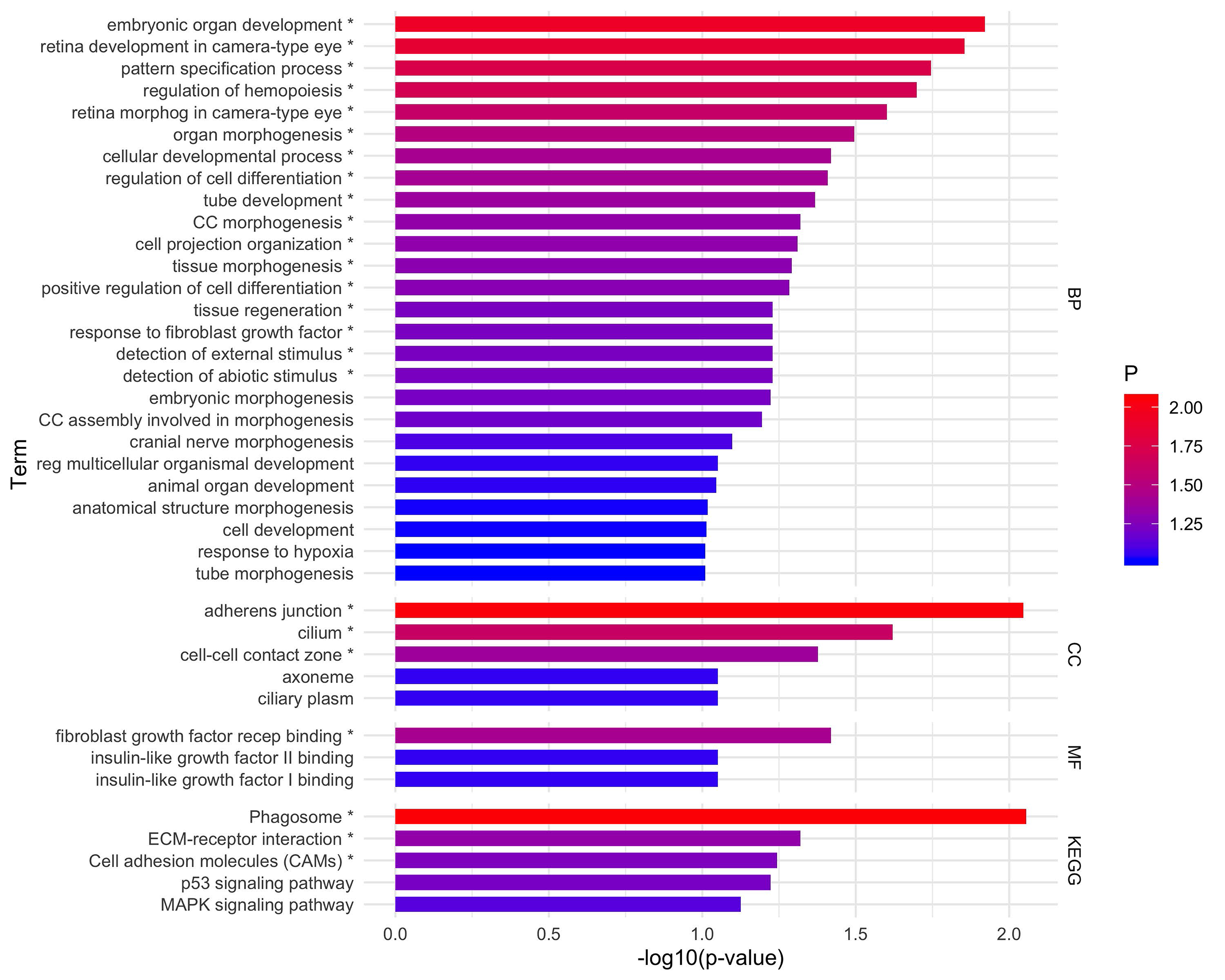
Enrichment analysis for GO and KEGG pathways term for strain C by DAVID. Each bar represents the -log10(p-value) for term. The colors represent the significance of each term; red color presents significant values and as the significance decreases the gradient turns blue. * significative terms.

## Discussion

Previous studies aiming at identifying selection signatures have been performed in different aquaculture species, including Atlantic salmon ^23,24,27^, brown trout ^25^ and Sockeye salmon ^26^. In Nile tilapia, there are only two studies of this kind and both have taken an inter-specific approach to detect signals of adaptation and selection in this species: The first one was carried out in the African cichlid lineages, including *O. niloticus* and other four representative species of cichlid family ^28^; and the second study was focused on *O. niloticus*, *O. mossambicus* and their hybrids ^29^. In this study, we evaluate the presence of selection signatures in three strains of Nile tilapia from two geographical origins: Brazil (strain A) and Costa Rica (strains B and C) using data from a whole-genome re-sequencing experiment and two complementary statistical methods (iHS and Rsb).

### Basic statistics and genetic structure

Genetic diversity (H_e_ and H_o_) was low and similar between all strains. The values of H_o_ were lower than values of H_e_, which suggests a decline of genetic diversity because of founder effect or effective population size (Yoshida et al. 2019). These results are similar to the values of H_e_ previously reported for the GIFT strain, which ranged from 0.2 to 0.4 ^29–31^. Other studies based on microsatellite data reported higher values for H_e_ that ranged from 0.6 to 0.7 for the GIFT strain ^4,32,33^. However, microsatellite markers typically exhibit a bias towards highly polymorphic loci and rapid mutation rates ^34^. A low genetic diversity is expected in domesticated populations, like the ones analyzed in this study given that, unlike their wild conspecifics, these populations can loss genetic diversity due to selective breeding and absence of gene flow with other populations ^35^.

Regarding the genetic structure, the PCA identified three clusters consistent with the three strains of Nile tilapia (Figure 1). The results of the ancestry analysis (Figure 2) accounted for the multiple origins of the strains (Best K = 6), and the similar genetic background of GIFT. GIFT is a synthetic strain composed of eight wild and farmed strains of Nile tilapia ^2,30^. According to Yoshida et al. (2019), these strains have been subjected to artificial selection for the improvement of growth-related traits in different geographic locations, which in conjunction to a founder effect and high admixture levels can be explaining the genetic structure of them.

The results reported in this study suggest a relatively low genetic diversity present in three different populations of farmed Nile tilapia, which can be explained by low effective population size (N_e_) and consequent genetic drift and inbreeding ^36^. Yoshida et al. (2019) reported N_e_ values of 159, 128, 78 for strains A, B and C, respectively. These values are expected as domesticated animals typically have values of N_e_<100 ^37^. Even though these values of N_e_ are small, they are enough to maintain inbreeding at acceptable rates and the necessary levels of diversity in the long-term for breeding populations ^38,39^.

#### Signatures of selection

Our results suggest that domestication and selective breeding have caused important changes in the genome of all the strains studied here. In all the analyses performed, selection signatures were often linked to genes involved in growth, reproduction and immune system, traits that are usually important for commercial purposes and subjected to direct or inadvertent selection.

The number of signals of selection across the genome detected by the Rsb test was higher than those detected by iHS in the three strains (Figure 3, Supplementary TableS1). Rsb test can identify selected alleles fixed or close to fixation ^40^, which could be indicative of older events of selection. Therefore, a higher number of selection signatures detected by Rsb might be associated with the first stages of domestication and the effect of artificial selection which may have fixed some favorable mutations in a given population ^41^. Instead, the iHS test has greater power when selected alleles are at intermediate frequencies ^42^. Hence, a lower number of regions detected by the iHS test could be accounting for more recent events of selection in these populations.

We focused the discussion on candidate genes underlying selection detected by both methods (Rsb and iHS) and which are potentially associated to the main biological processes involved in selection and adaptation to captive environment which were over-represented in the gene annotation (e.g. growth, immune response and reproduction). For instance, we found several genes associated with growth-related traits, including as ATOH8, CSNK1A1, COPS6, LARP7, COL25A1, RIN2, collagen alpha-2 (VI) chain, DIS3L2, MYO18A. The ATOH8 gene may regulate the function of the transcriptional complex (GATA and FOG) that play a role in the development of multiple organ systems in mammals and fish ^43^. CSNK1A1 gene products are involved in the canonical Wnt signaling pathway, which in turn regulates the development and differentiation of multiple tissues ^44^. We also found two genes associated with the collagen family, COL6A2 and COL25A1 gene. The first one have a role during the muscle development in zebrafish ^45^. The second one corresponds to a molecule that is required for the fusion of myoblasts into myofibers ^46^. We detected three genes (LARP7, MYO18A, and COPS6) associated with growth which were previously identified in zebrafish. LARP7 gene was previously associated with a critical role in cardiomyocyte proliferation and response to injury ^47^. MYO18A gene likely functions in the adhesion process that maintains the stable attachment of myofibers to the extracellular matrix and muscle integrity during early development ^48^. COPS6 gene plays different roles in early embryonic development, including dorsoventral patterning, convergent extension movement, and brain formation ^49^. We found two genes which were previously identified to be associated with growth in sheep: DIS3L2 gene was associated with body weight and height ^50,51^ and RIN2 gene was associated with growth and meat production ^52^.

Regarding genes involved in immune response, we found molecules such as Ladderlectin, BTN1A1, Mucin-2-Like, and Cadherin-1 (CDH1) among others. More specifically, the *Ladderlectin* gene has been previously associated with an innate immune response mechanism, which corresponds to plasma pattern recognition for bacterial, fungal and viral hemorrhagic septicemia virus in rainbow trout ^53^. The *BTN1A1* gene was associated with immune homeostasis as it participates in processes such as T cell selection, differentiation, and cell fate determination ^54^. The *Mucin-2-Like* gene was found to be up-regulated in susceptible catfish challenged against *Flavobacterium columnare* ^55^, bacterium which can also replicate in tilapia mucus ^56^.

We found the *Borealin-2* gene, which likely plays an important role during oogenesis and early embryogenesis in rainbow trout ^57^. Finally, associated with the adaptation to environmental stimulus, we also found the 60S Ribosomal Protein L34 gene (RPL34), which has been found to be up-regulated in white muscle tissues following thermal stress in bluefin tuna (*Thunnus orientalis*) ^58^.

#### Functional enrichment analysis

Based on the functional enrichment analysis, it was possible to detect biological processes that could be under the effect of domestication and directional selection in these strains of Nile tilapia. Most of the GO and KEGG pathways terms detected were associated with traits relevant in farmed populations. Therefore, here we focus on GO and KEGG pathways terms that could be associated with the process of domestication and artificial selection (Supplementary Table S4). These terms were associated with: growth (G), immune system (I), reproduction (R), behavior (B) adaptation to the environment (A) and nervous system (N). We detected 25 of 54 (Table 3), 26 of 90 (Table 4) and 39 of 67 (Table 5) enriched terms, in strains A, B and C, respectively.

These results are consistent with the expected effect of domestication and adaptation in a culture system in fish, as well as the effect of efficient directional selection for growth rate in these fish populations. Aquaculture systems are characterized by less complexity than natural conditions. Thus, they tend to decrease adaptive pressures for many traits and induce selective pressures for other traits (Lorenzen et al. 2012). For example, farmed animals generally present changes in their life history patterns, redirecting their resources toward processes like growth and reproduction (Lorenzen et al. 2012). In this study the most frequent term (GO and KEGG) was related with growth-related traits for all strains, representing 78% of all terms (70 terms: 19, 20 and 31 in strains A, B and C, respectively). This result could be due to the fact that the genetic improvement of the base synthetic strain (GIFT) and all of the derived strains studied here, was based on the improvement of the growth (Eknath and Hulata 2009). For instance, Latin American strains (A, B, and C) have been improved for growth-related traits for about ten generations. Similarly to Hong Xia et al. (2015), we identified the Wnt canonical signaling pathway in strains A, B and C, which has been linked to the regulation of tissue development and differentiation ^59^. Finally, these findings account for the possible polygenic nature of the growth trait ^60^, i.e., the growth of fish is controlled by large numbers of small effect genes ^61^. Polygenic dependence is suggested as the growth term was found independently in GO and KEGG pathways terms.

We found only 4% of enriched terms associated with reproduction in strains A and C. This result might be suggesting a trade-off between growth and reproductive traits in these strains. In fish, resources are generally prioritized for growth until maturity^62^. For instance, Bolivar et al. (1993) analyzed growth and reproduction traits in seven strains of Nile tilapia (three wild and four domesticated strains) and found three phenotypic categories of females based on age at first spawning: early spawn, late spawn, virgin females. Females with phenotypes “late spawn” and “virgin females” showed the same growth performance as males ^63^.

Through captivity fish populations present changes in behavior traits as well; including aggressiveness, which often increases, and other traits such as foraging, anti-predator and reproductive behavior, which frequently decrease in complexity ^36^. In strain A, GO terms related to behavior trait such as, cognition (GO:0050890, associated to activities with thinking, learning, and memory) and learning (GO:0007612, linked to adaptive behavioral change occurs as the result of experience) were found. In strains B and C, we detected terms associated with adaptation to the environment: Detection of external/abiotic stimulus (GO:0009581, GO:0009582), Melanogenesis (dre04916), Response to hypoxia (GO:0001666) and P53 signaling pathway (dre04115) (Supplementary Table S4). These characteristics might be representing advantageous adaptations for farming systems ^64^.

## Conclusion

In this study we detected several genomic regions putatively underlying selection in three farmed populations of Nile tilapia. These regions harbor interesting candidate genes, which may be associated with the adaptive processes to captivity and traits of economic importance which have been subjected to artificial directional selection. In addition, the result of the enrichment analysis of all candidate genes identified was often linked to production traits; most commonly to growth, accounting for the efficient genetic improvement for growth-related traits in these three strains. Our results may be relevant for a better understanding of genes underlying traits of interest in aquaculture and the effect of domestication in the genome of Nile tilapia.

## Methods

### Fish samples

A total of 326 individuals of farmed Nile tilapia from three commercial strains cultivated in two different countries of Latin America were included in this study (Table 1). Strain A was originally imported from Malaysia to Brazil in 2005, and samples for this study were obtained from the breeding population of AquaAmerica, Brazil. This strain is derived from the GIFT strain, a mixture of four Asian domestic strains from Israel, Singapore, Taiwan and Thailand with four wild populations from Egypt, Senegal, Kenya, and Ghana (Eknath and Hulata 2009). Strain B and C were introduced from the Philippines (station Carmen aquafarm) to Costa Rica in 2005, and samples were obtained from the Aquacorporacion Internacional (Costa Rica) breeding population. Strain B is a mixture of an eight-generation GIFT strain, two wild populations from Egypt and Kenya and fish from Strain C, which in turn originated from a mixture of Asian domestic strains from Israel, Singapore, Taiwan and Thailand. The sampling protocol were performed accordance with *Comité de Bioética Animal, Facultad de Ciencias Veterinarias y Pecuarias, Universidad de Chile, Chile* (certificate Nº19179-VET-UCH).

### Sequence data and quality control

DNA from all individuals was purified from fin-clip samples using a Wizard Genomic DNA purification kit (Promega). The DNA libraries were prepared and sequenced by Illumina HiSeq 2500 (Illumina, USA) as described by Cáceres et al. (2019) and Yáñez et al. (2019). Reads were aligned to the Nile tilapia reference genome (O_niloticus_UMD1, https://www.ncbi.nlm.nih.gov/bioproject/PRJNA344471, Conte et al. (2017)) with BWA MEM ^68^. The discovery of variants was made with the Genome Analysis Toolkit (GATK) software (https://www.broadinstitute.org/gatk/) ^69^. The variant coordinates were updated to the latest version of the genome (O_niloticus_UMD_NMBU, Conte et al. (2019)).

The variants were filtered using the VCFtools software (version 0.1.15) ^71^ and SNPs that did not pass the following quality control (QC) were removed: (1) indels, (2) SNPs with more than two alleles, (3) Quality of phred score < 30, (4) call rate < 90% from SNPs, (5) mitochondrial SNP, (6) Deviated from Hardy-Weinberg Equilibrium with Bonferroni correction (HWE, p-value<1×10-9), and (7) minor allele frequency (MAF) < 0.05. Step 6 and 7 were applied on each strain separately. The individuals exhibiting variant call rate below 80% were removed. Closely related individuals may bias estimates of allelic and haplotypic frequencies, and thus might mask signature of selection ^19^. To avoid highly related individuals between samples we performed an analysis of identical by descent (IBD) with PLINK v1.09 ^72^, where one individual from pairs of animals with high values of IBD were excluded (Details in Table 2). We imputed missing genotypes and inferred haplotypes using BEAGLE v.3 ^73^. with default parameters. Finally, the effect of each variant over the potential gene function was predicted using SnpEff ^74^.

### Basic statistics and population structure analysis

Genetic diversity among populations was calculated through observed and expected heterozygosities (Ho and He) using PLINK v1.09. We measured genetic differentiation among strains using pairwise Weir and Cockerham’s Fst estimator in StAMPP package of R ^75^.

To examine genetic structure among populations, we first performed a principal component analysis (PCA) implemented PLINK v1.09 (Chang et al. 2015). Second, to infer the number of populations between strains we used the maximum likelihood analysis of individual ancestries by ADMIXTURE software ^76^. The number of ancestral populations (K) was set from 1 to 10 and the optimal K was selected based on the lowest cross-validation error and a visual inspection of co-ancestry values.

### Signatures of selection

We used two complementary methods to detect signatures of selection. Both methods are based in extended haplotype homozygosity (EHH), that correspond to the probability that two randomly chosen chromosomes carrying the core haplotype are identical by descent ^77,78^. The first method is the intra-population standardized integrated haplotype score (iHS) ^42^; the second is the inter-population standardized log-ratio of integrated EHH (iES) between pairs of populations (Rsb) ^40^. Both methods were applied using REHH package ^77^.

iHS compares EHH between alleles within one population, i.e. the area under the curve of the derived and ancestral alleles ^78^. This method requires the information of ancestral allele identification for each SNP. We defined ancestral allele as the one with the highest frequency ^77^. Standardized iHS was defined as:

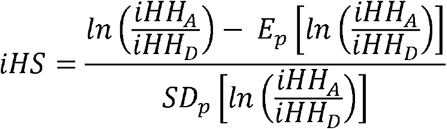

Where iHHa and iHHd corresponded to integrated EHH score for ancestral (A) and derived (D) core alleles respectively. The iHS values were calculated separately within-populations (A, B and C). Positive or negative values of iHS indicate that haplotype carrying the ancestral or derived allele respectively present an extended haplotype (unusually high haplotype homozygosity).

Rsb compares EHH profiles of the same allele between pairs of populations ^40^. This method was defined as the natural logarithm of the ratio between iESpop1 and iESpop2, where iES represent the integrated EHHS (site-specific EHH) for both alleles of each SNP within each population. Rsb was calculated between populations (AB, AC, and BC). This method requires no information of ancestral and derived alleles. Positive values of Rsb indicate iESpop1 is greater than iESpop2, i.e., pop1 has longer haplotype than pop2, therefore suggest positive selection in the alternative population (pop1) ^24^. Conversely, negative values score suggests positive selection in a reference population (pop2) ^24^.

Identifying the causal variant at a site of selection is hard, but if single nucleotide polymorphisms (SNPs) on a selected haplotype are closely linked to a candidate gene, this information could be used as evidence of a potential sign of selection near that gene ^14^. Therefore, candidate regions for selection were defined as those genomic positions containing SNPs with values of iHS and Rsb above the threshold. The threshold used to set the significance of iHS and Rsb methods corresponds to 7.4 (-log10(p-value), with Bonferroni correction). We used a range of 500kb upstream and downstream to the candidate SNP to explore for candidate genes under selection. The genes intersecting these regions were considered a candidate to selection and detected using BEDTools ^79^.

### Functional enrichment analysis

Using all candidate genes under selection, detected by both methods, we performed a BLAST against zebrafish (*Danio rerio*) proteins, using the genome reference annotations from NCBI of both species. An enrichment analysis was conducted using the online tool David Bioinformatics platform ^80,81^ to detect GO and KEGG pathways terms.

## Availability of data and material

The datasets generated during and/or analysed during the current study are available the online digital repository Figshare, access number https://figshare.com/s/4118213db62982996ed6.

## Acknowledgments

The authors are grateful to Aquacorporación Internacional and AquaAmerica for providing the Nile tilapia samples. Doctoral fellowship CONICYT (21171369). BIRDS MSCA RISE 2015 project.

## Authors’ contributions

MIC performed the analysis and wrote the initial version of the manuscript. MEL, DD contribute with analysis, discussion, and writing. GC performed DNA extraction. GY, GC and DGU reviewed manuscript. JMY, MIC, and MEL conceived, designed the study and writing. All authors have reviewed and approved the manuscript.

## Ethics approval and consent to participate

Nile tilapia sampling procedures were approved by the Comité de Bioética Animal from the Facultad de Ciencias Veterinarias y Pecuarias, Universidad de Chile (certificate Nº19179-VET-UCH).

## Consent for publication

Not applicable

## Supplementary information

Supplementary Table S1. SNPs detected by iHS and Rsb by chromosome in each strain.

Supplementary Table S2. List of all genes detected by iHS methods.

Supplementary Table S3. List of all genes detected by Rsb methods.

Supplementary Table S4. List of terms and definitions of enrichment analysis.

